## Supplementary material for "Cellular heterogeneity–adjusted clonal methylation (CHALM) provides better prediction of gene expression"

### Cell heterogeneity adjusted clonal methylation better predicts gene expression

#### Supplementary Tables

Supplementary Table 1: List of promoter CGIs

Supplementary Table 2: List of differentially methylated promoter CGIs between lung tumor-normal pair.

Supplementary Table 3: List of de novo differentially methylated regions between LUAD and SCLC.

Supplementary Table 4: List of datasets used in this study

### Supplementary Figures

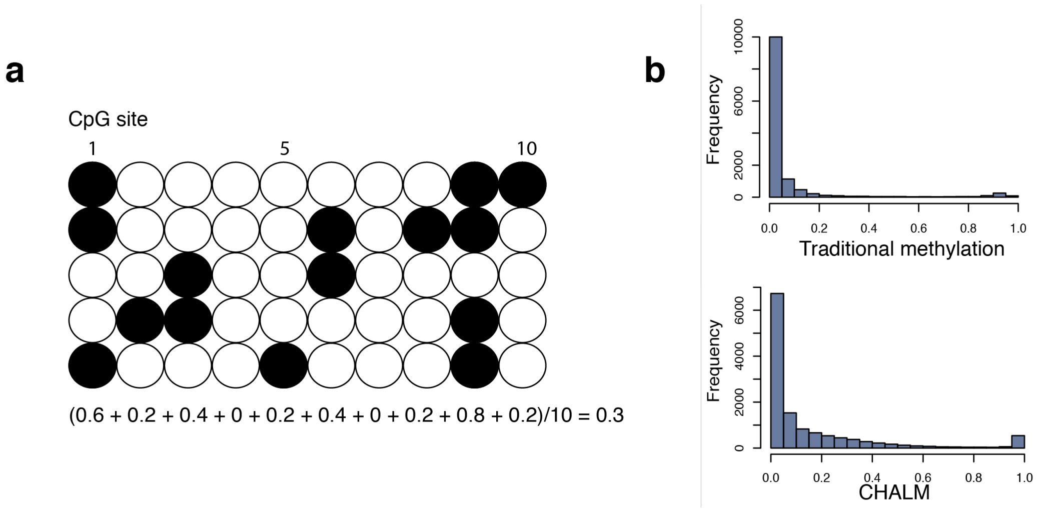


**Supplementary Figure 1. Extended method description. (a)** Plot illustrating the traditional method for quantifying the methylation level of a promoter CGI. **(b)** Histogram of traditional methylation and CHALM methylation level of promoter CGIs.

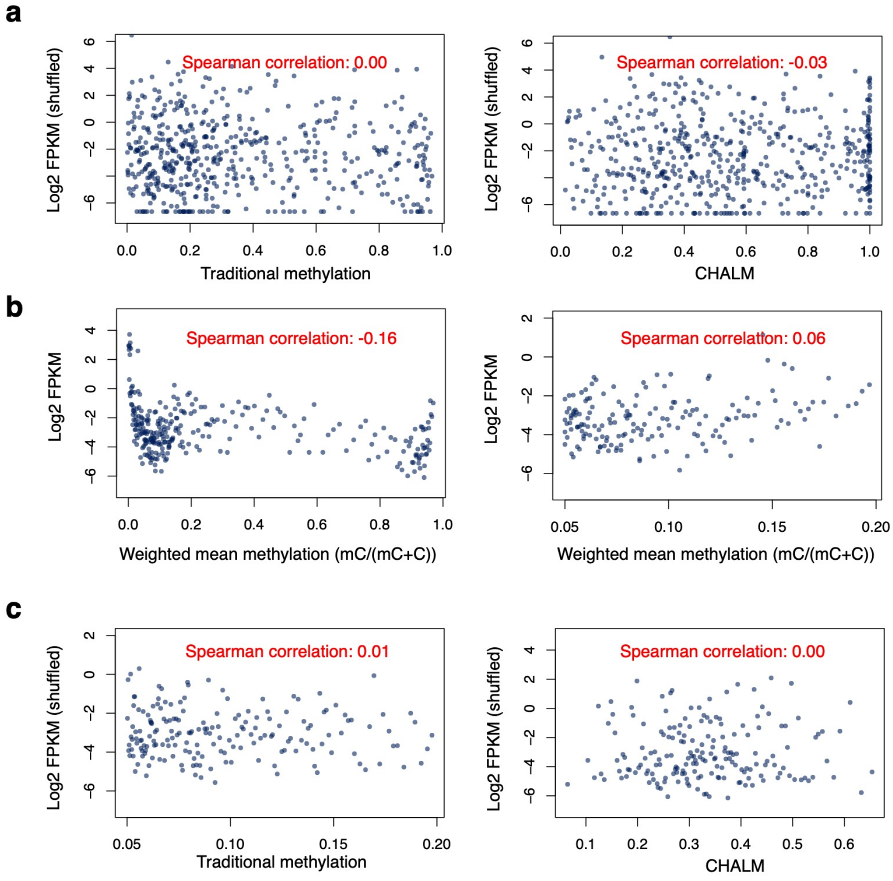


**Supplementary Figure 2. More on correlation between DNA methylation and gene expression. (a)** There is no correlation between DNA methylation and shuffled gene expression on a genome-wide scale. **(b)** The correlation between another traditional method, i.e. "weighted mean methylation level", and gene expression is similar to what is observed for mean methylation level, which is the traditional method mainly discussed in this work. Left: balanced promoter CGIs set. Right: low methylated promoter CGIs. **(c)** There is no correlation between DNA methylation and shuffled gene expression for low methylated genes.

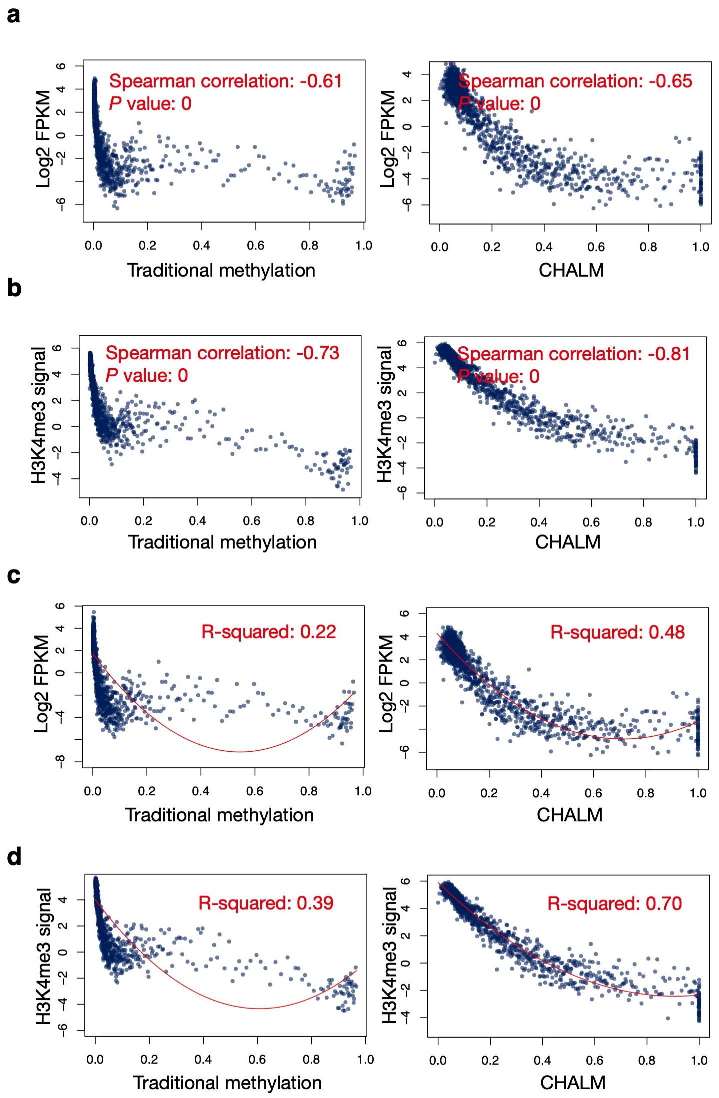


**Supplementary Figure 3. CHALM better predicts gene expression and H3K4me3 level (CD3 primary cell dataset).** All promoter CGIs are used for plotting the scatter plots showing the relationship between methylation level and gene expression **(a)** or H3K4me3 level **(b)**. Polynomial regression is used to fit the relationship between methylation level and gene expression **(c)** or H3K4me3 level **(d)**. The R-squared values are calculated with the raw data of all promoter CGIs.

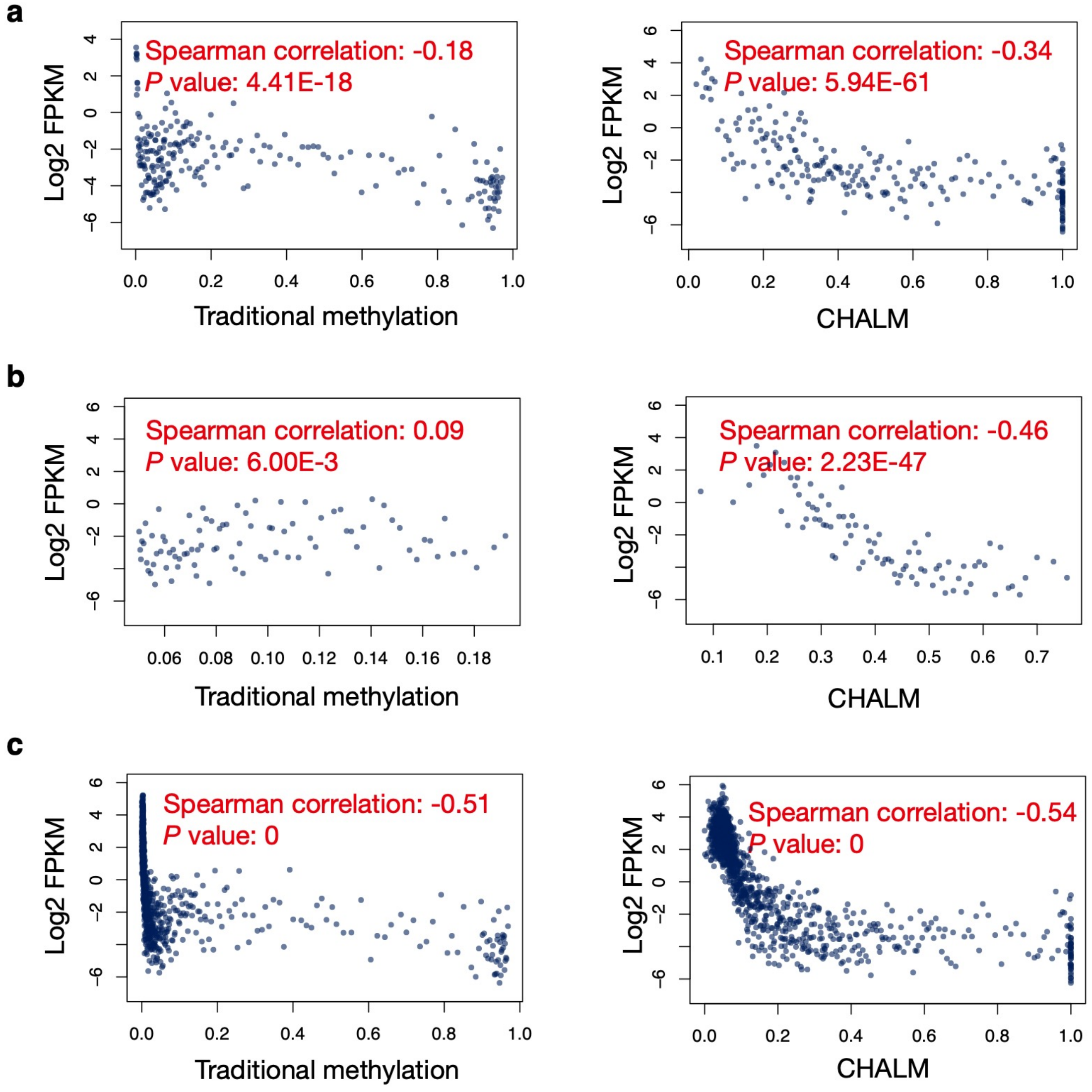


**Supplementary Figure 4. CHALM better predicts gene expression (CD14 primary cell dataset).** Balanced promoter CGIs set **(a)**, low methylated promoter CGIs **(b)** and all promoter CGIs **(c)** are used for scatter plots. Significant correlation comparison permutation *P* value can be observed in **(a)** and **(b)**.

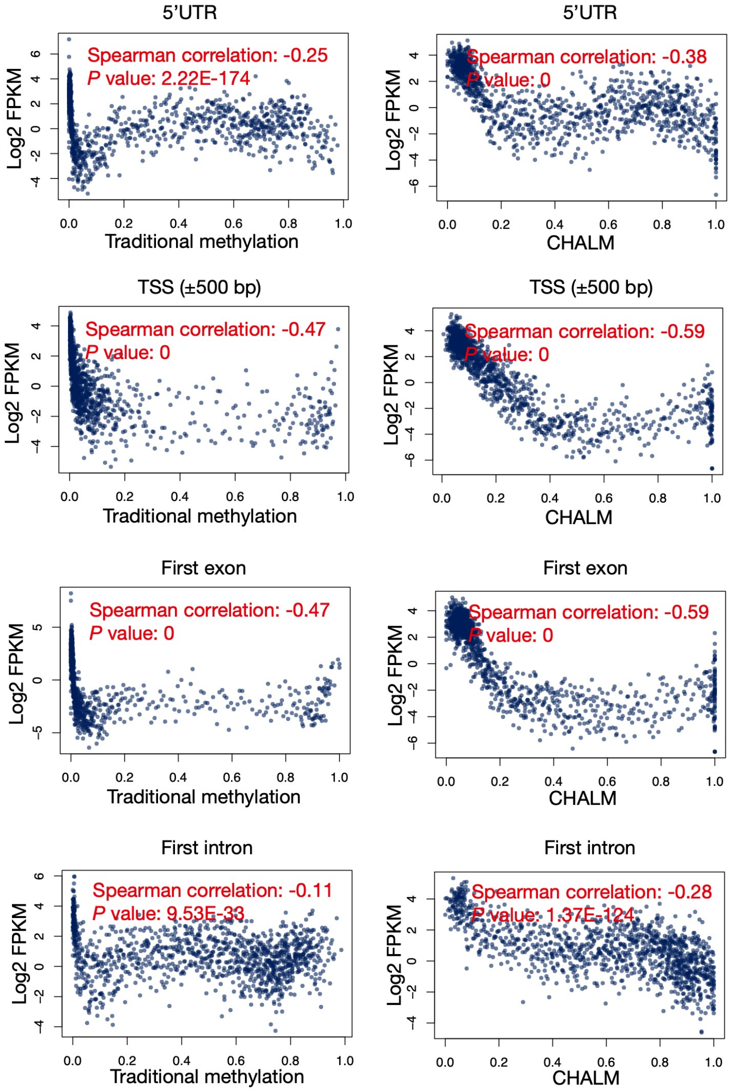


**Supplementary Figure 5. Correlation of methylation level and gene expression in 5'UTR, TSS surrounding region, first exon and first intron.** Left panel: Traditional methylation. Right panel: CHALM. CD3 primary cell is used for plotting. Correlation comparison (between traditional method and CHALM) permutation *P* values for all four regions are less than 0.05.

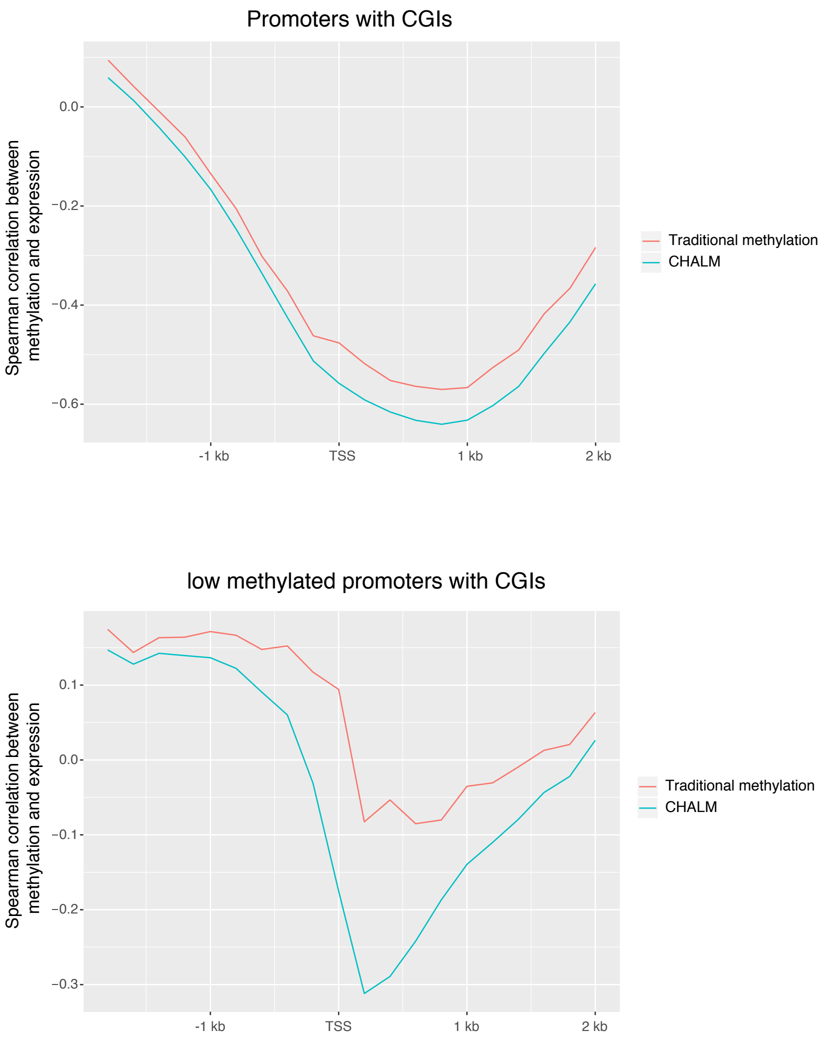


**Supplementary Figure 6. The correlation of CHALM and traditional methylation with gene expression in the TSS surrounding region.** Top panel: all promoters with CGIs are included for analysis. Bottom panel: low methylated promoters with CGIs are included for analysis.

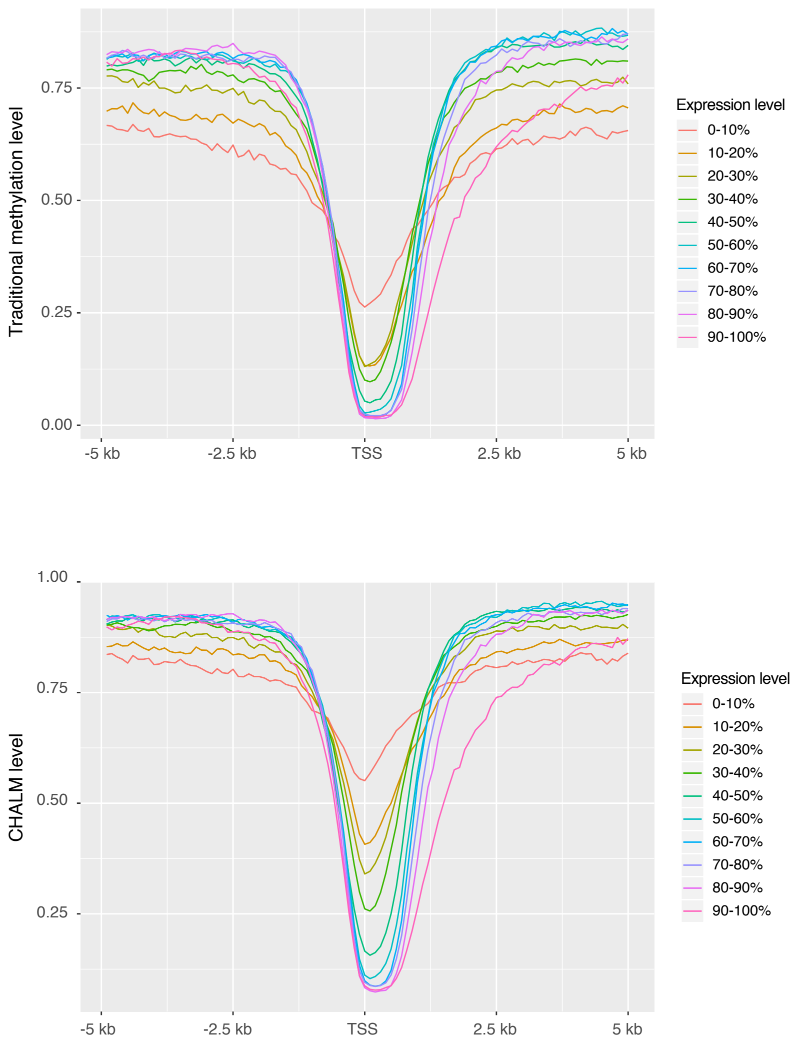


**Supplementary Figure 7. Metagene plots show that CHALM is better correlated to gene expression.** Top panel: Traditional methylation. Bottom panel: CHALM. CD3 primary cell is used for plotting.

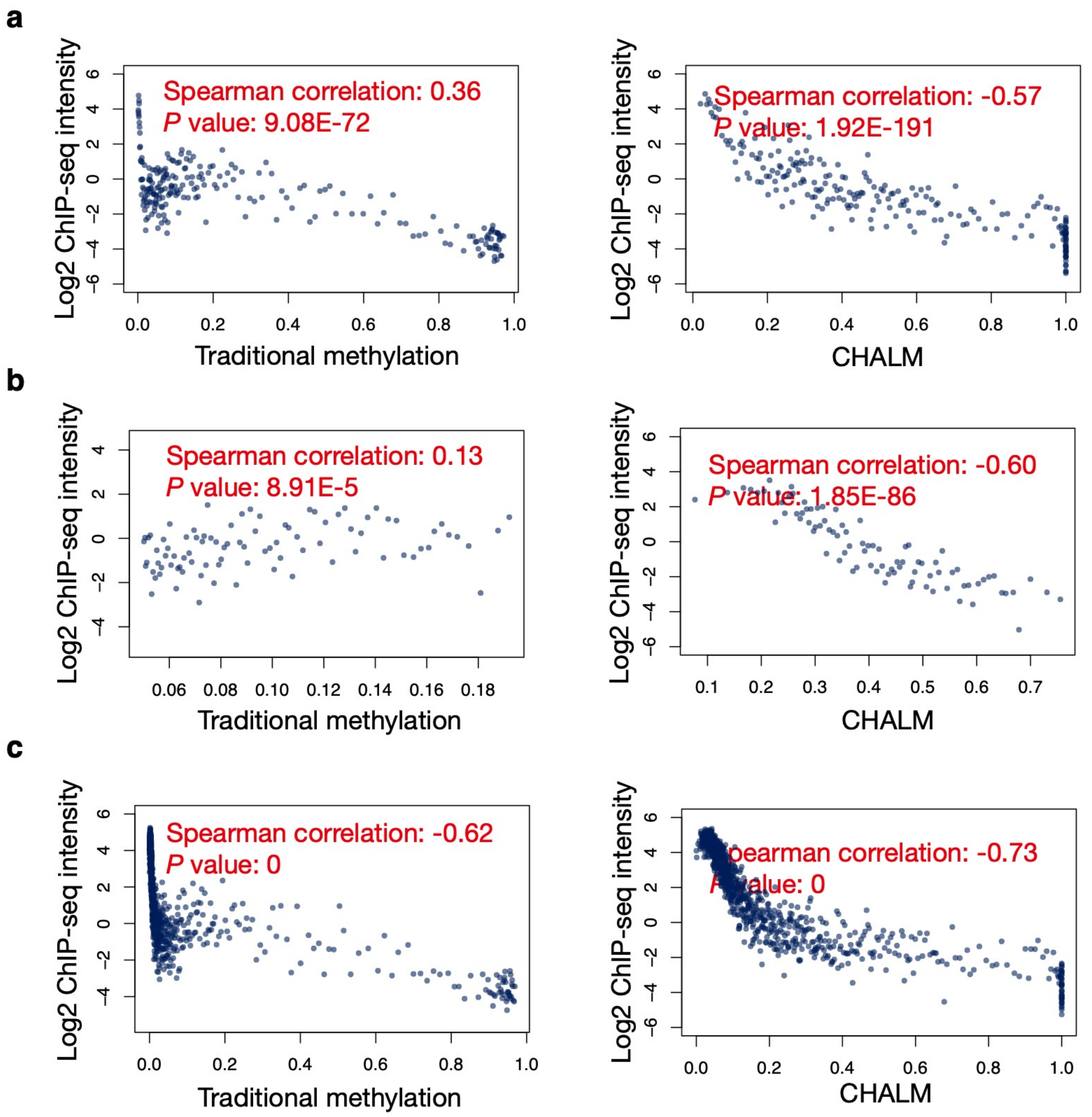


**Supplementary Figure 8. CHALM better predicts H3K4me3 level (CD14 primary cell dataset).** Balanced promoter CGIs set **(a)**, low methylated promoter CGIs **(b)** and all promoter CGIs **(c)** are used for scatter plots. Significant correlation comparison permutation *P* value can be observed in **(a)**, **(b)** and **(c)**.

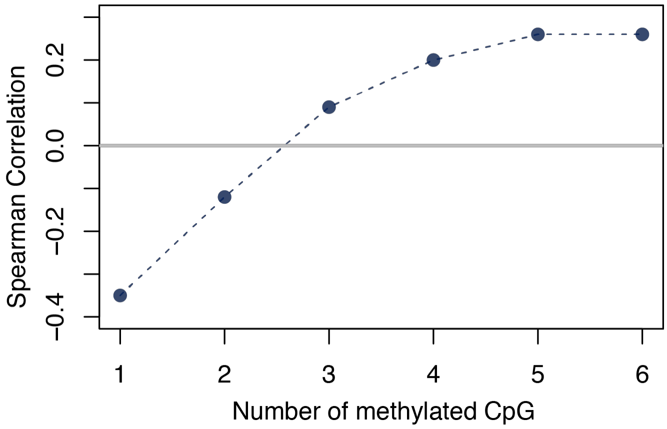


**Supplementary Figure 9. CHALM performs best when methylated read has least one mCpG site.** The definition of the methylated read is the sequencing read with at least N mCpG. The X-axis represents different values of N. The Y-axis represents the spearman correlation between gene expression and CHALM of low methylated genes under given N value.

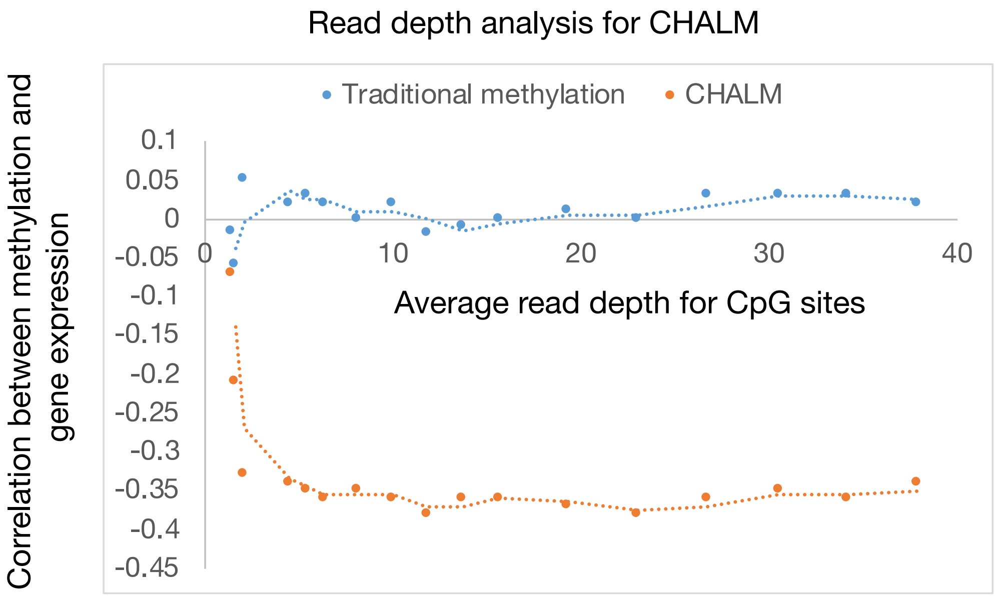


**Supplementary Figure 10. The influence of sequencing depth on CHALM performance.**

X-axis shows the average CpG depth in the promoter CGIs. Y-axis shows the correlation of the methylation level to the gene expression for the low methylated promoter CGIs.

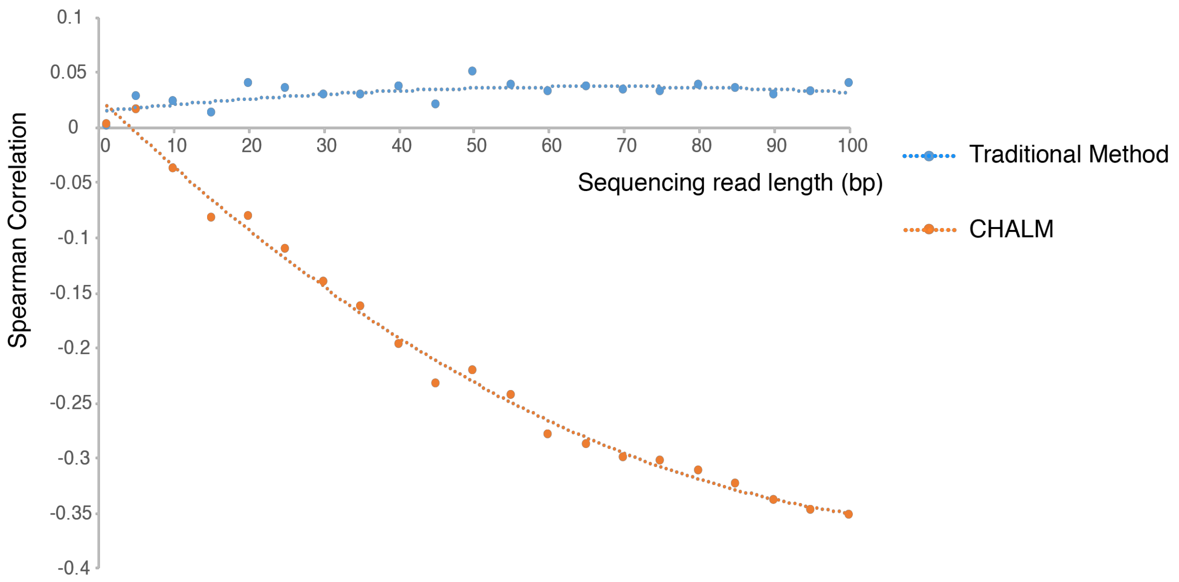


**Supplementary Figure 11. CHALM performs bad for short-read dataset.** The scatter plot showing the influence of sequencing read length to the CHALM performance. The X-axis represents the sequencing read length. The Y-axis represents the spearman correlation between gene expression and CHALM of low methylated genes under given read length. For simplicity, CHALM with single-end mode is used.

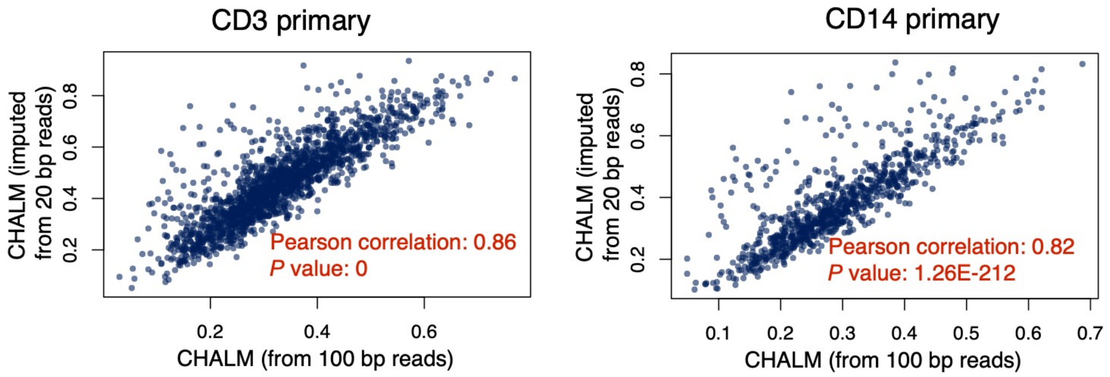


**Supplementary Figure 12. The SVM imputation method can impute the CHALM value with high accuracy.** Scatter plots showing the high correlation between the imputed CHALM values and the measured CHALM values in CD3 primary dataset (left) and CD14 primary dataset (right).

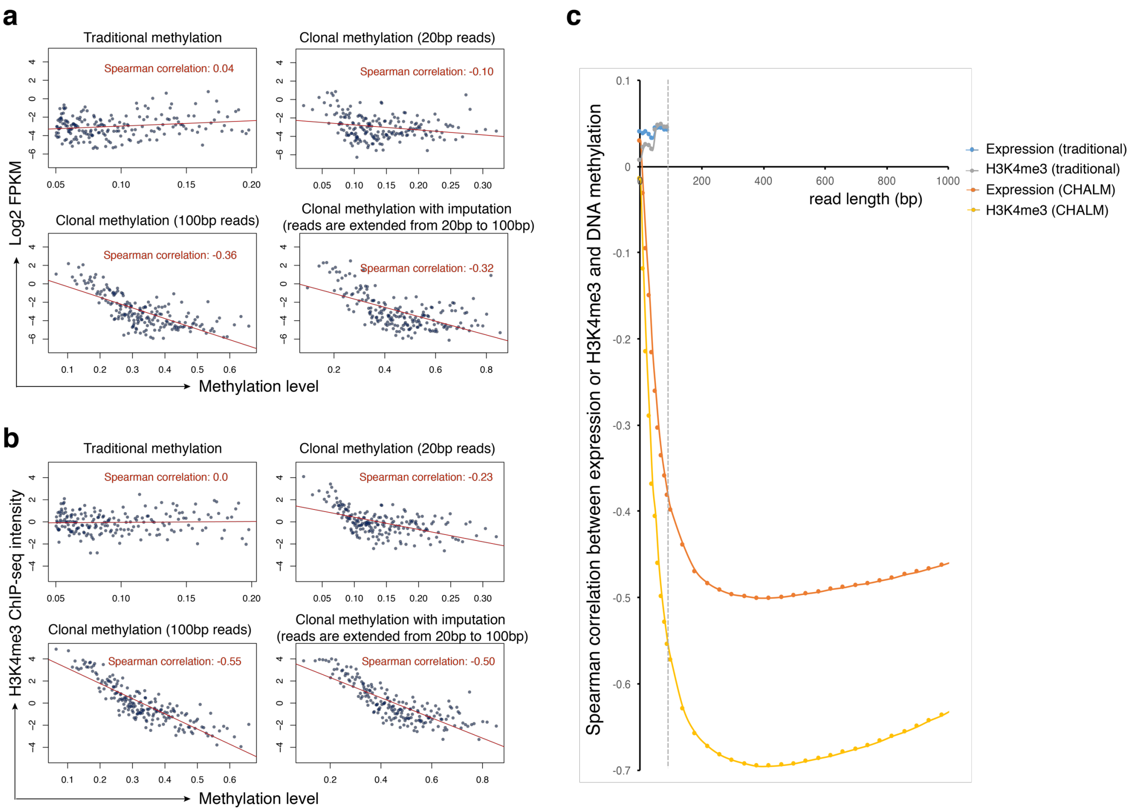


**Supplementary Figure 13. Extend sequencing read length by imputation. (a)** Validate the imputation method by the correlation between gene expression and DNA methylation of low methylated genes. The sequencing read length of original dataset is 100 bp. The sequencing read is sheared into 20 bp long to generate an artificial short-read dataset. Imputation method is then used to extend the 20 bp read of the short-read dataset back into 100bp to generate an imputation dataset. The spearman correlation between gene expression and CHALM is calculated for original dataset, short-read dataset and imputation dataset. **(b)** Validate the imputation method by the correlation between H3K4me3 and DNA methylation level of low methylated genes. **(c)** Extend the 100 bp sequencing read to generate artificial long-read datasets. The correlation between expression or H3K4me3 and CHALM for low methylated genes is then calculated for these long-read datasets. The X-axis represents the read length and the Y-axis represents the spearman correlation of the low methylated genes. For simplicity, CHALM with single-end mode is used.

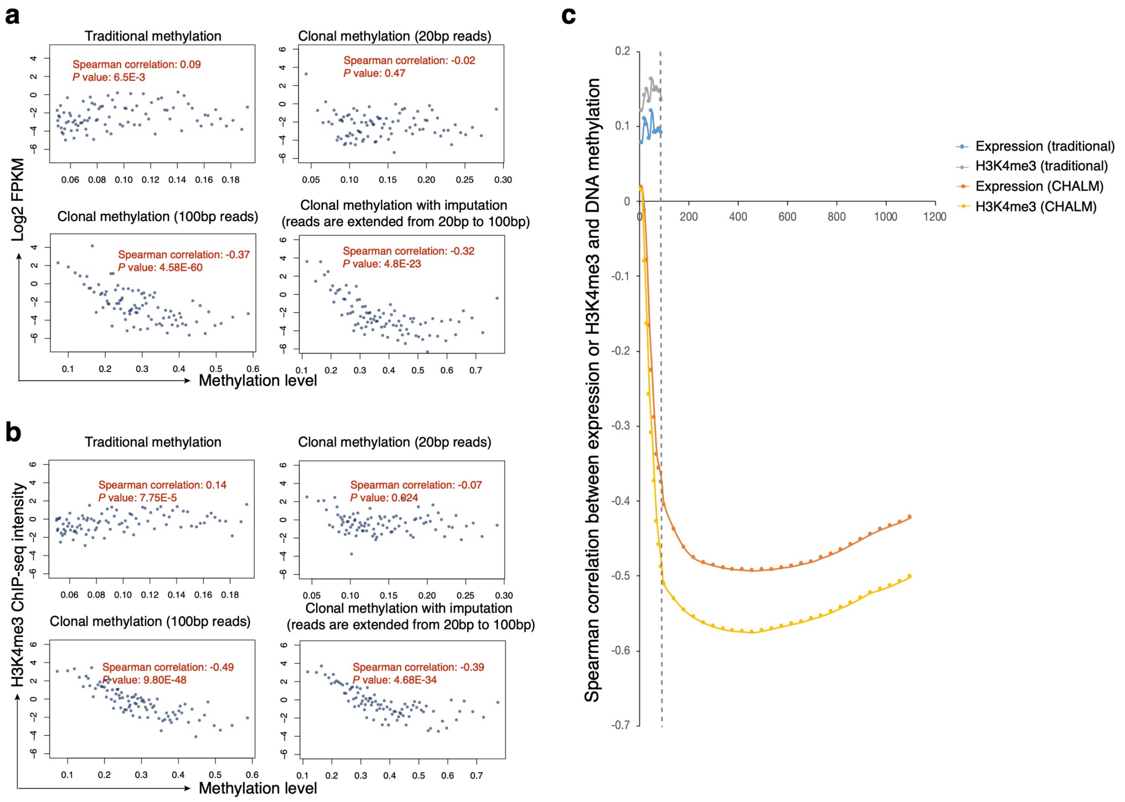


**Supplementary Figure 14. Extend sequencing read length by imputation (CD14 primary cell dataset).**

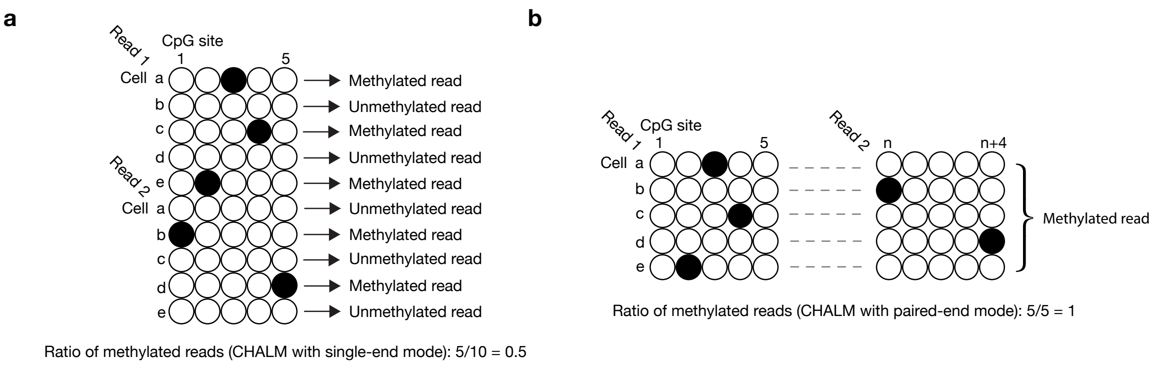


**Supplementary Figure 15. Single-end mode and paired-end mode of CHALM. (a)** For paired-end data, CHALM with single-end mode will treat a paired reads as two independent single-end reads. **(b)** CHALM with paired-end mode will consider the paired reads as a single long read (concatenated by read 1, insert region and read 2). If any CpG site is methylated in read 1 or read 2, the whole long read will be considered as methylated read.

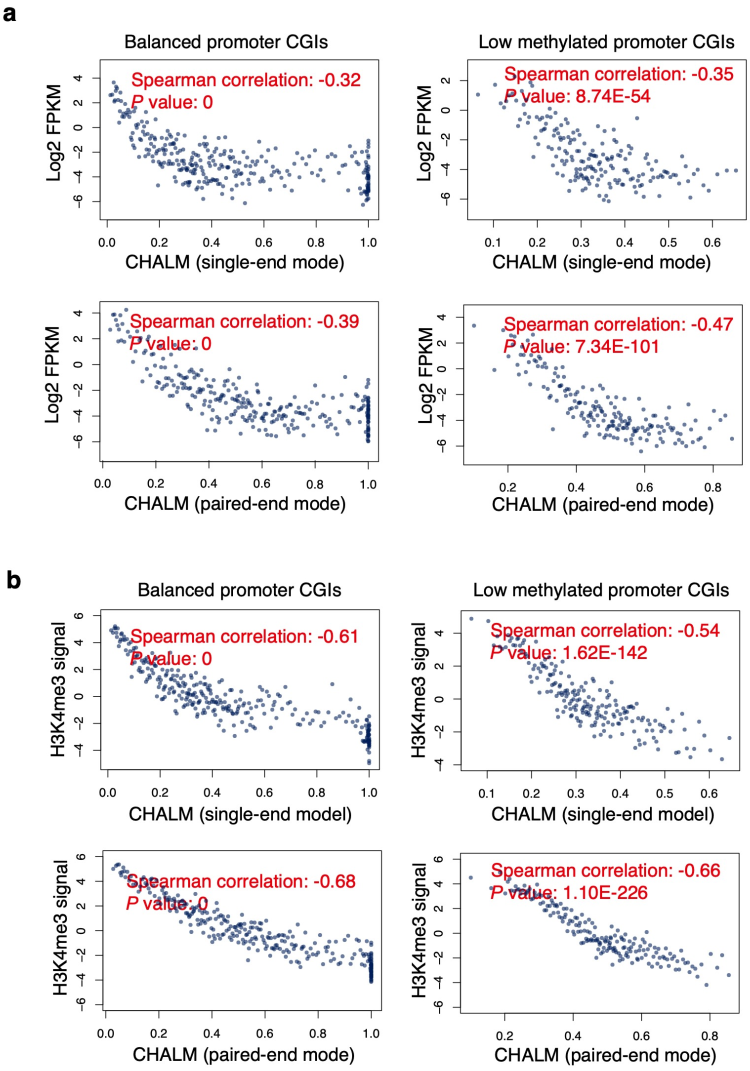


**Supplementary Figure 16. Compare the performance of single-end mode and paired-end mode of CHALM.** Correlation of CHALM to the gene expression **(a)** and H3K4me3 level **(b)** in the promoter CGIs.

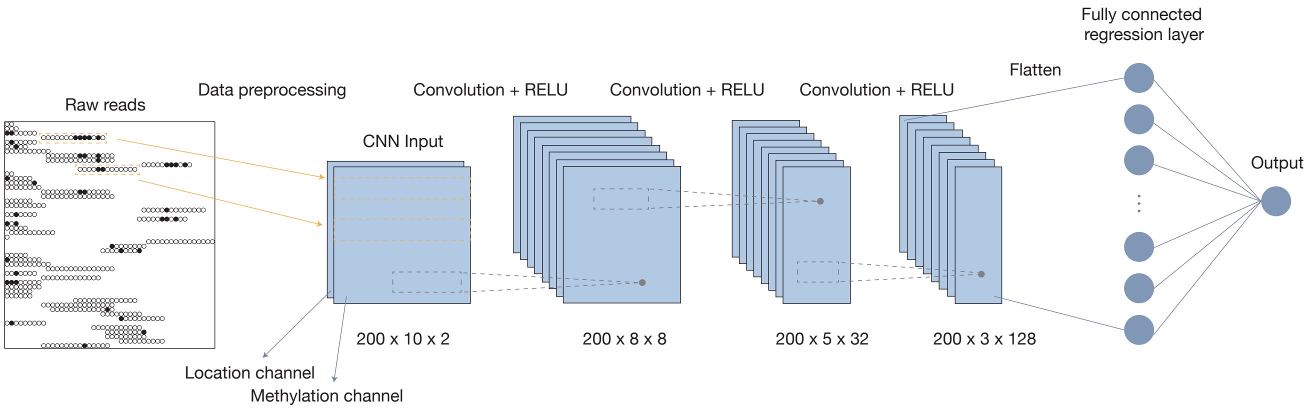


**Supplementary Figure 17. Deeping learning prediction framework.** Raw WGBS sequencing reads mapped to a promoter CGI region are processed into an image-like data structure, which has two channels for containing CpG methylation status and the read's distance to the transcription start site. Each row represents one single sequencing read. The image-like data structure is first scanned by different 2D filters for convolution. After three convolution layers and one fully connected layer, a final linear regression layer is used for gene expression prediction.

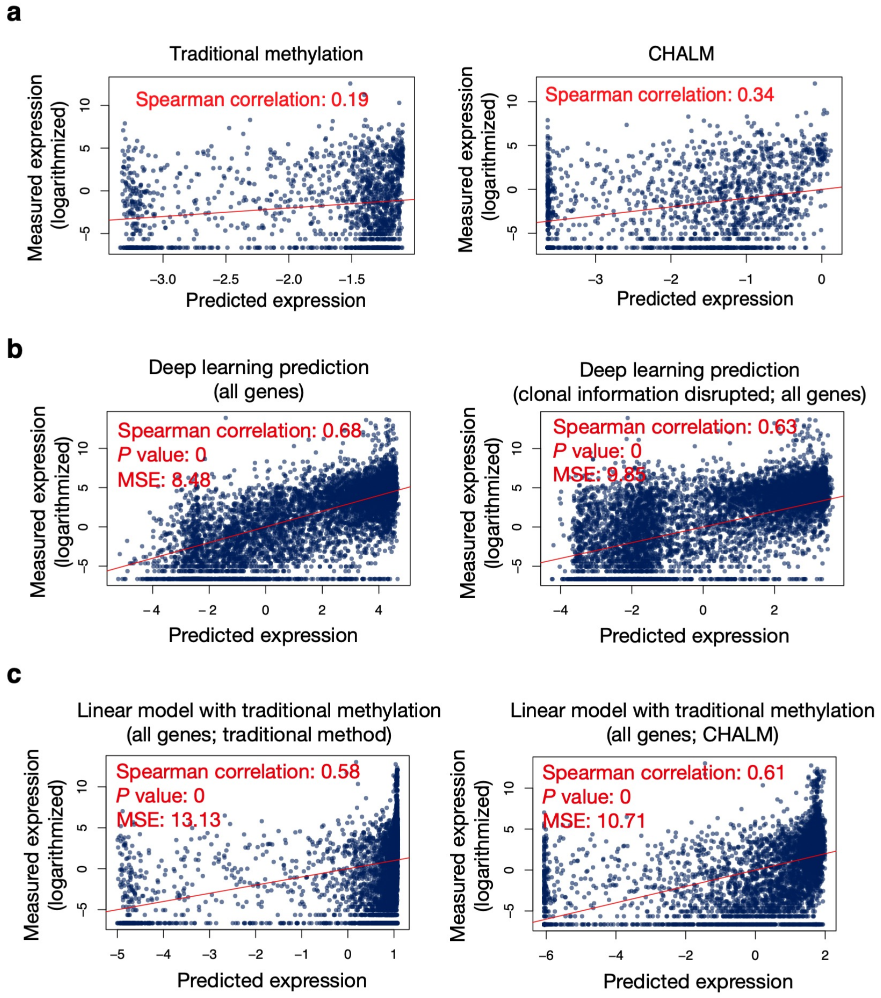


**Supplementary Figure 18. Gene expression prediction by traditional methylation or CHALM methylation level (CD3 primary cell dataset). (a)** Linear regression model is used for gene expression prediction with the balanced promoter CGIs set. Correlation comparison permutation *P* value: < $1 \times 10^{4}$. CNN model **(b)** and linear model **(c)** are used for gene expression prediction with all promoter CGIs. For simplicity, CHALM with single-end mode is used.

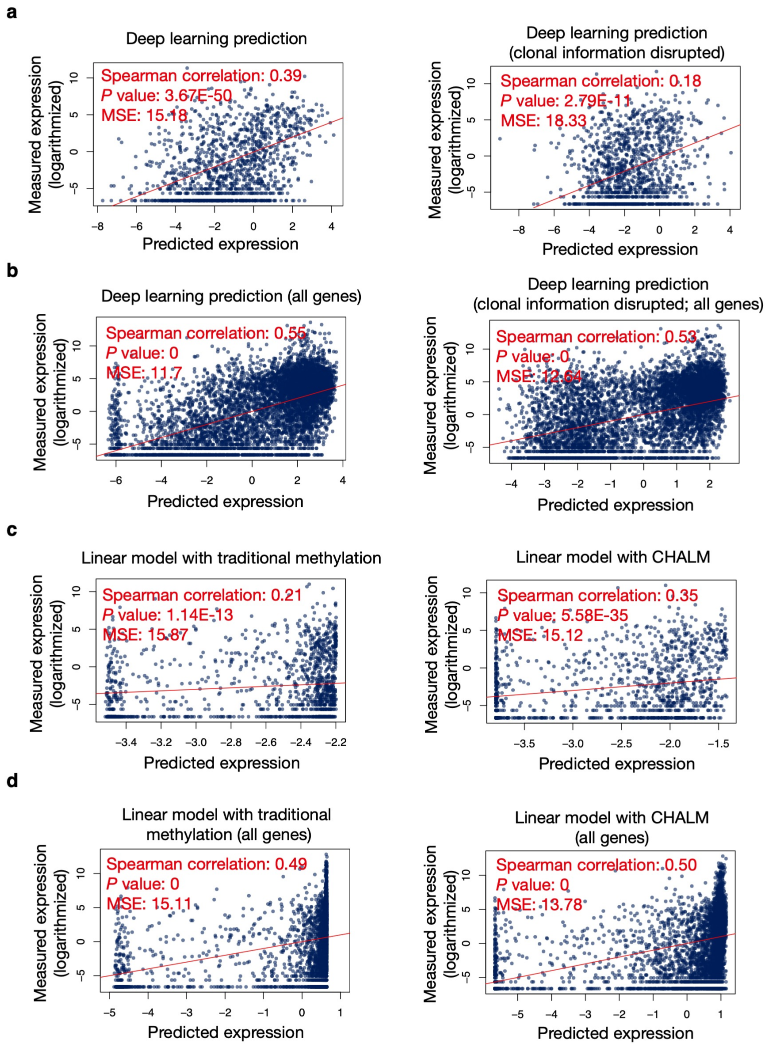


**Supplementary Figure 19. Deep learning model using raw sequencing read as input for predicting gene expression (CD14 primary cell dataset).** Significant correlation comparison permutation *P* value can be observed in **(a)** and **(c)**.

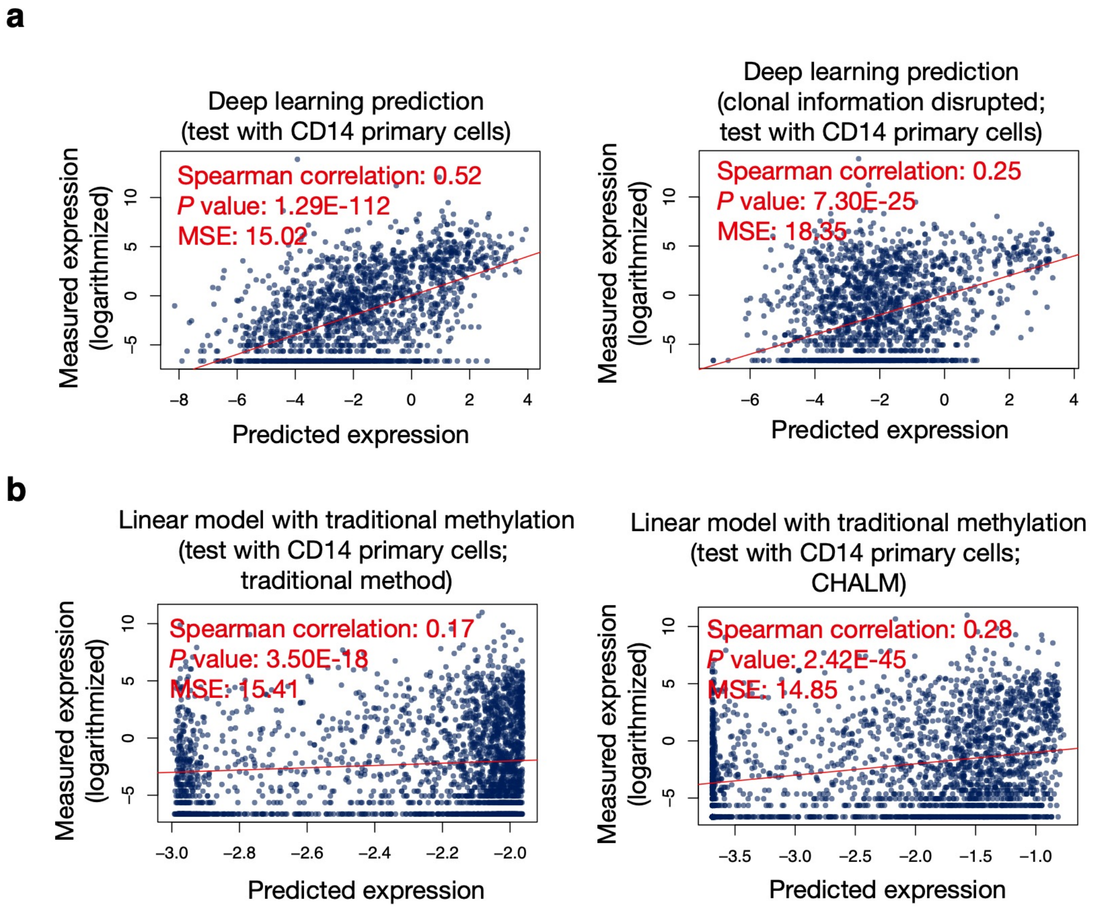


**Supplementary Figure 20. Deep learning model validation by independent dataset.** The CNN model **(a)** and linear model **(b)** are trained with CD3 primary cell dataset but tested by the CD14 primary cell dataset. Correlation comparison permutation *P* value is less than 1x$10^{4}$ in **(a)** and **(b)**.

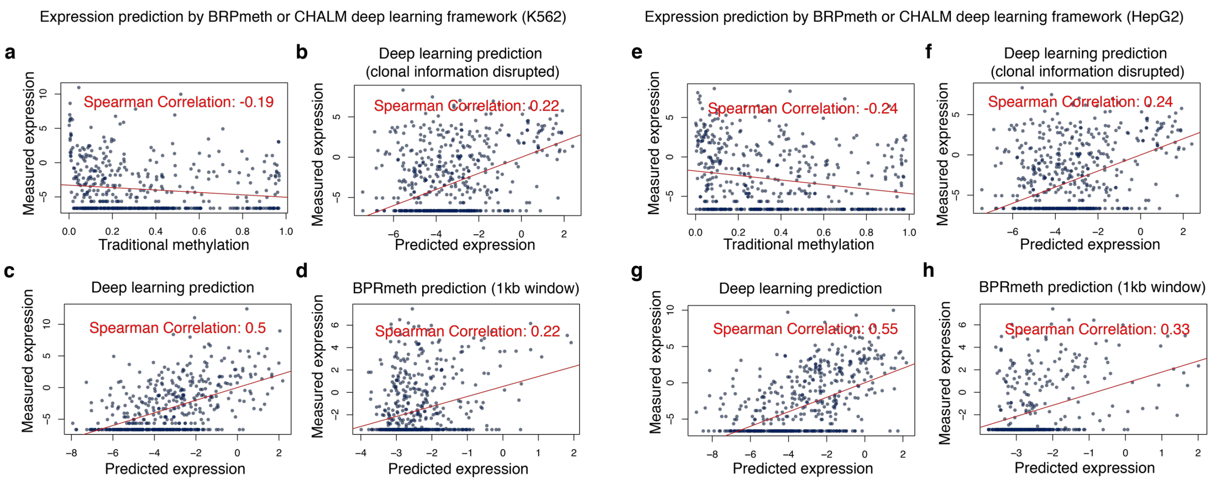


**Supplementary Figure 21. Gene expression prediction by BPRmeth or CHALM deep learning framework for K562 and HepG2. (a)** or **(e)** Scatter plot shows the spearman correlation between traditional methylation and gene expression of promoter CGIs in test datasets. **(b)** or **(f)** The prediction results of CHALM deep learning framework when the clonal information is disrupted. **(c)** or **(g)** The prediction results of CHALM deep learning framework. **(d)** or **(h)** The prediction results of BPRmeth. The window size is set as 1kb, which is about the average size of promoter CGIs.

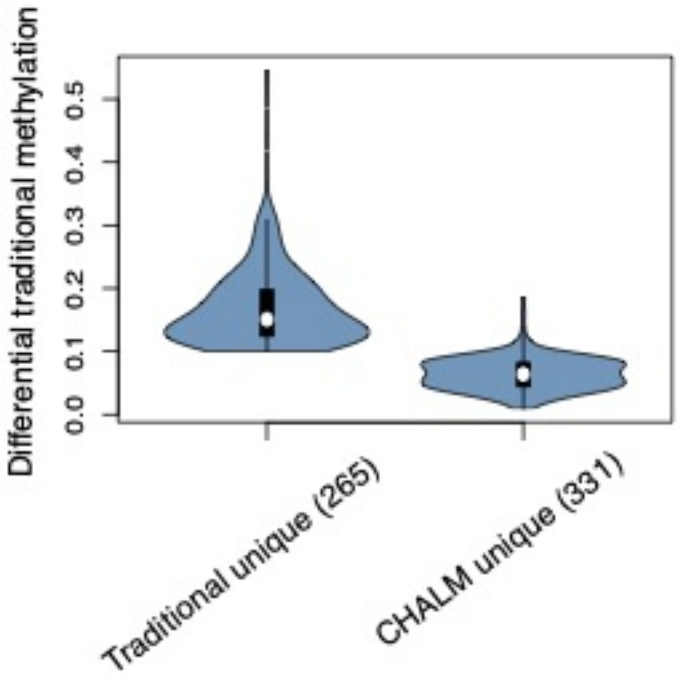


**Supplementary Figure 22. CHALM unique hypermethylated genes have minimal traditional methylation change.** Violin plot shows the differential traditional methylation change of hypermethylated gene sets unique to traditional method or CHALM.

**a**

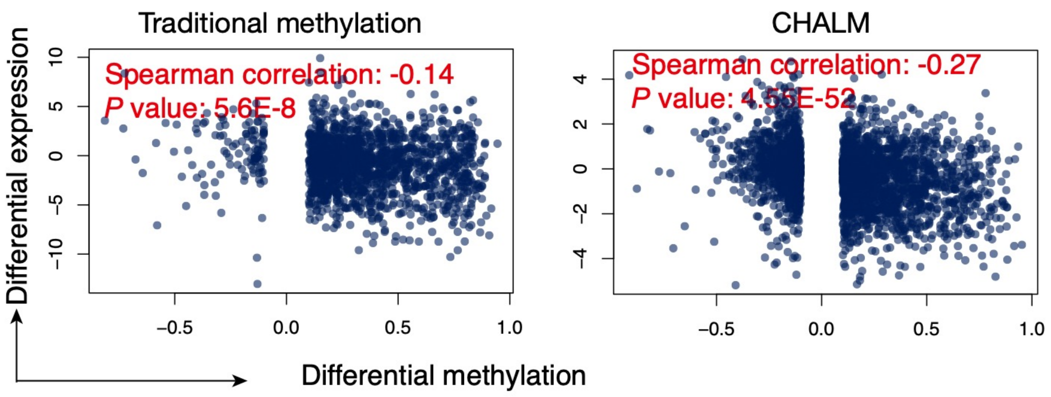


**b**

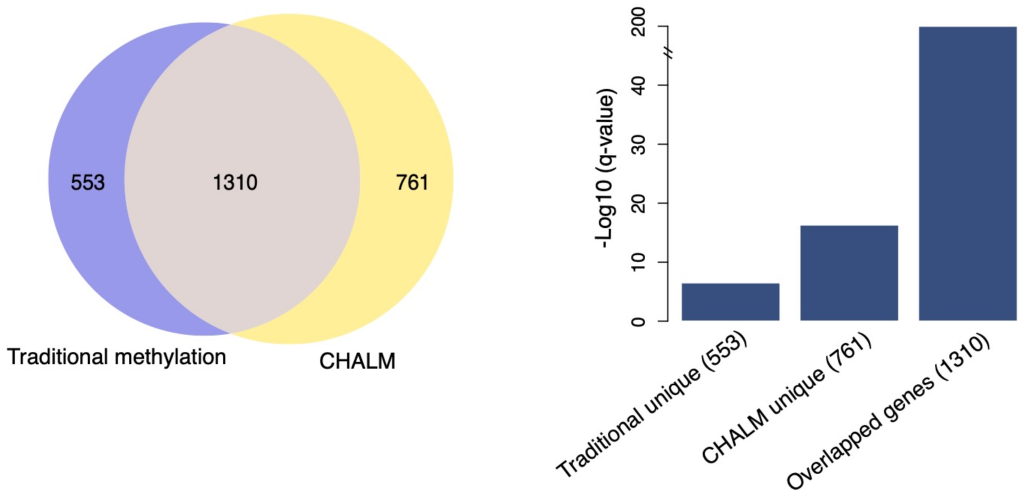


**Supplementary Figure 23. The CHALM methods provides better identification of hypermethylated promoter CGIs during tumorigenesis of uterine corpus endometrial carcinoma (UCEC). (a)** Scatter plots show the correlation between differential expression and differential methylation calculated by the traditional and CHALM methods. Promoter CGIs with a significant methylation change between normal and UCEC tumor tissue were plotted. Correlation comparison (between traditional method and CHALM) permutation *P* value: $< 1 \times 10^{-4}$. **(b)** A large fraction of hypermethylated promoter CGIs identified by the traditional method can be recovered using the CHALM method, as indicated by the Venn diagram. Bar plot shows enrichment of the H3K27me3 peak in three different gene sets.

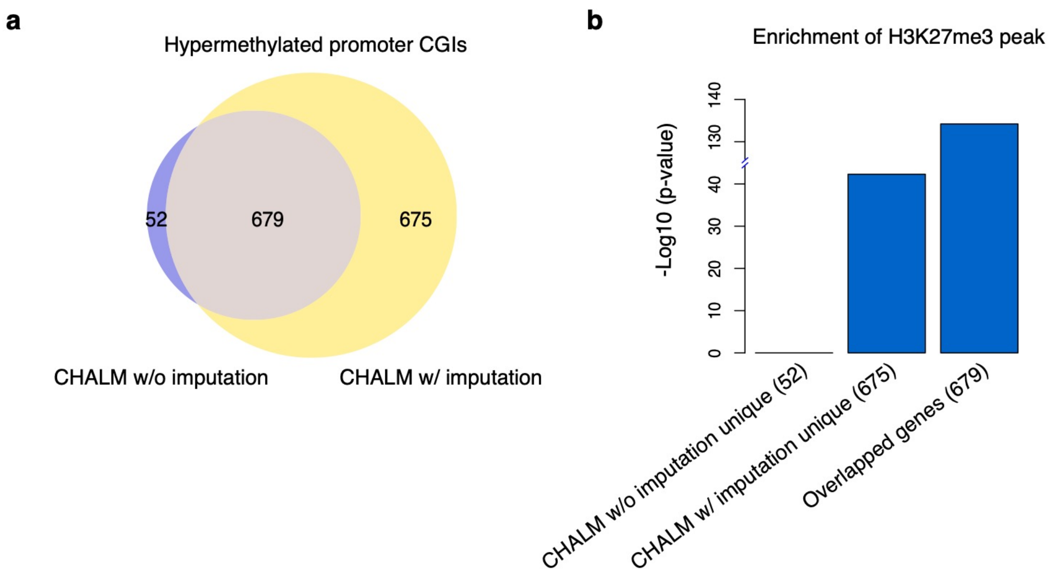


**Supplementary Figure 24. CHALM with SVD imputation identifies more hypermethylated promoter CGIs in the lung cancer dataset. (a)** Venn diagram showing that the CHALM of imputed long-read datasets is capable of identifying more hypermethylated promoter CGIs in lung normal-tumor tissue comparison. **(b)** The enrichment of H3K27me3 peak for different gene sets in **(a)**.

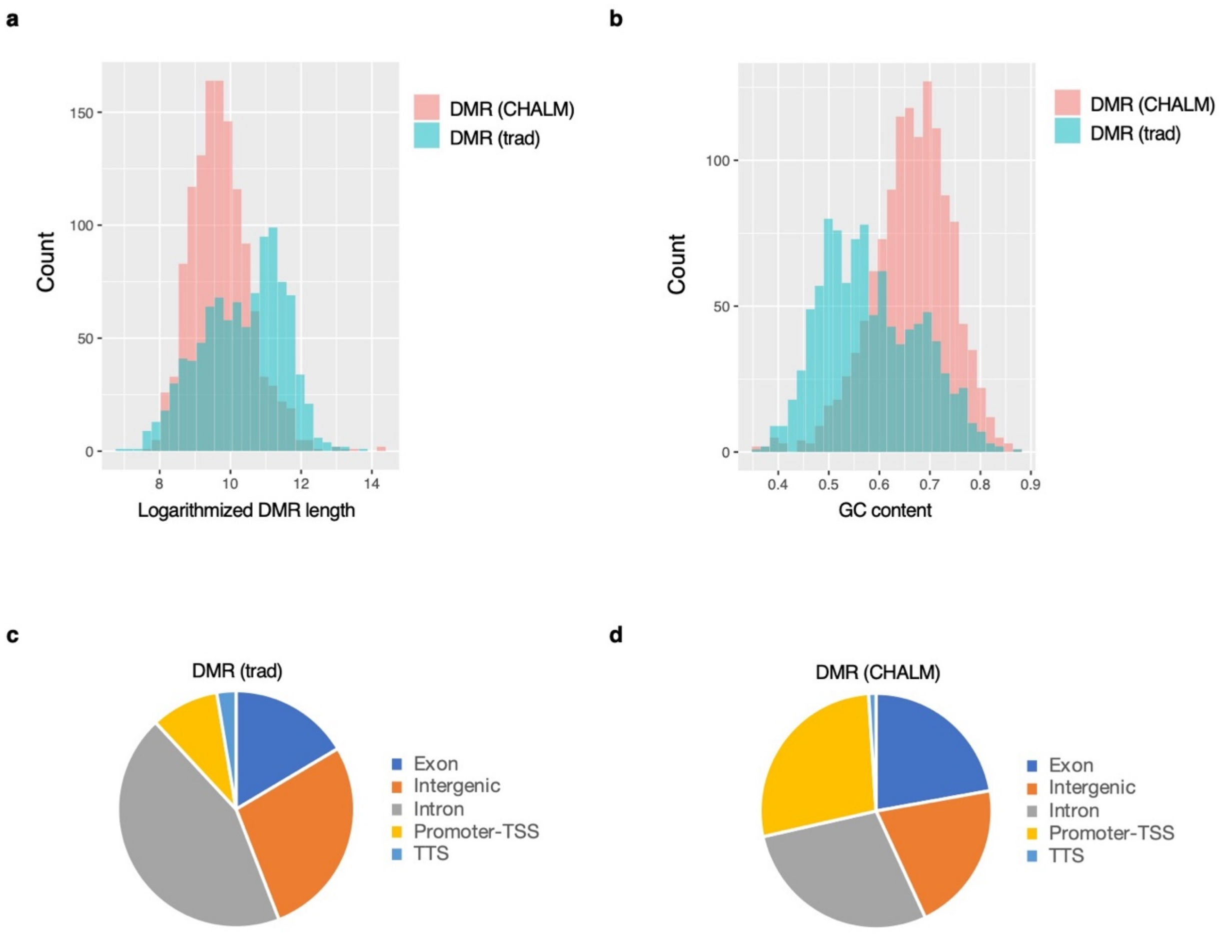


**Supplementary Figure 25. Comparison of the unique DMRs identified by traditional methylation and CHALM.** DMR length **(a)** and GC content **(b)** of the unique DMRs identified by traditional method and CHALM. Genomic annotation of traditional unique **(c)** and CHALM unique **(d)** DMRs.

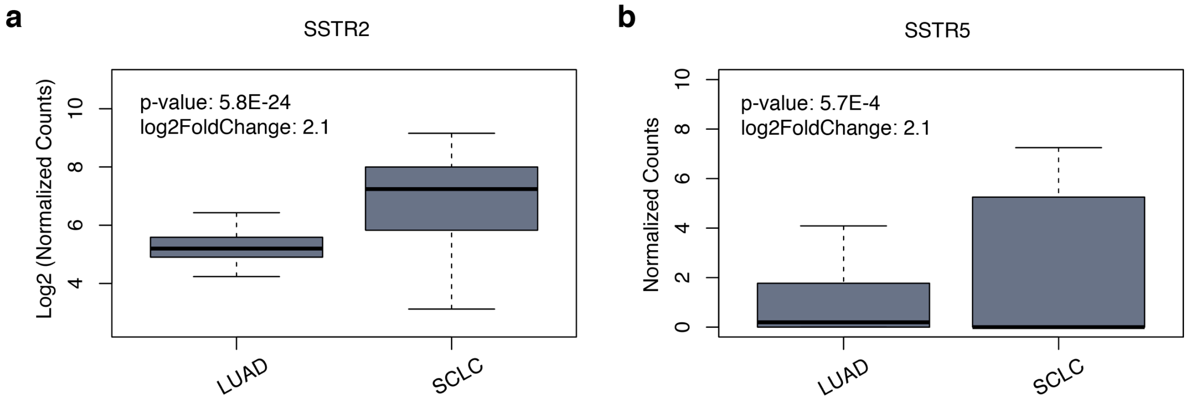


**Supplementary Figure 26. Expression of SSTR2 and SSTR5 in LUAD and SCLC patients.** 79 LUAD and SCLC patients are used. For SSTR5, the normalized count is not logarithmically transformed since this gene is low expressed.

**a**

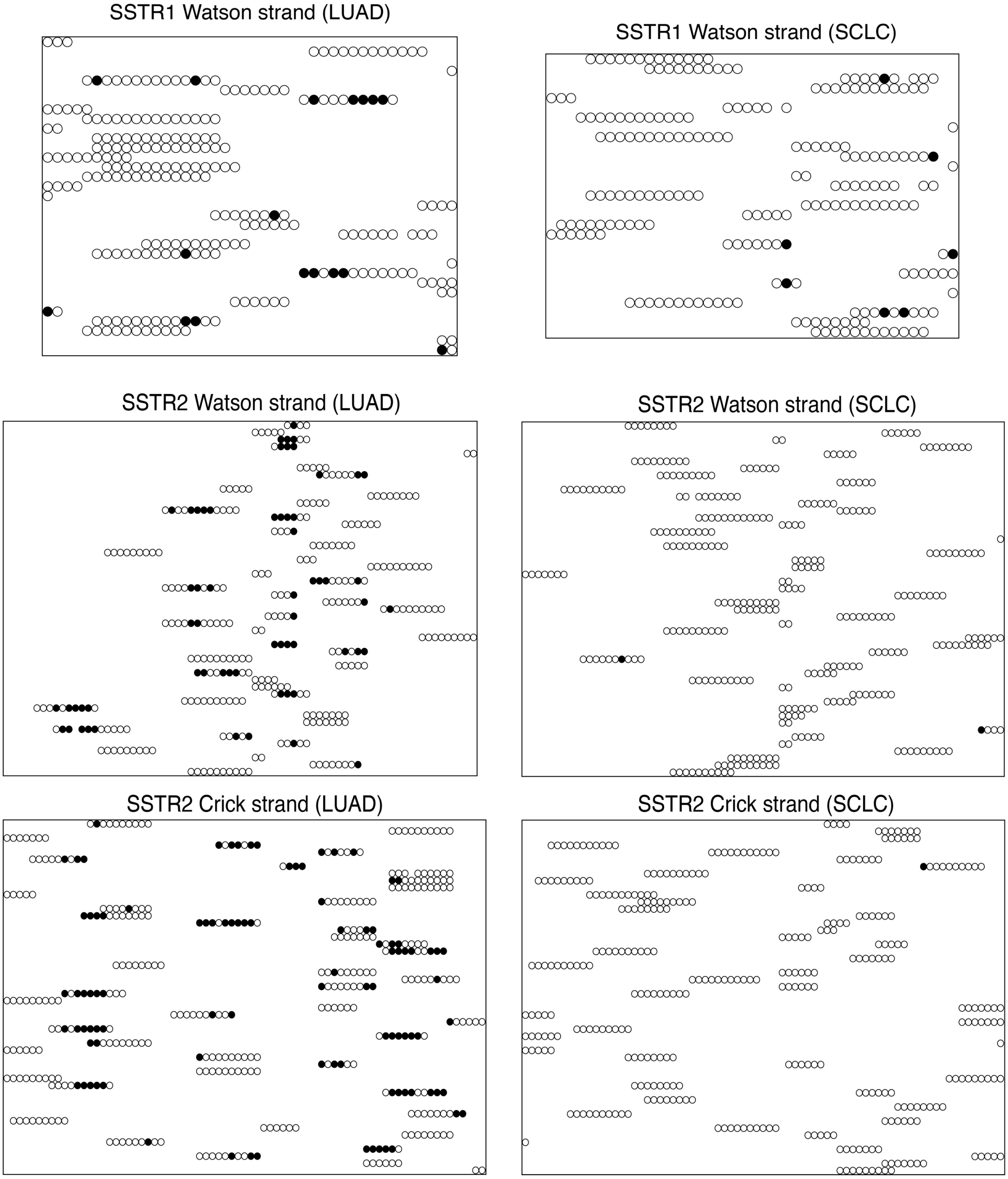
SSTR1 Watson strand (LUAD)
SSTR1 Watson strand (SCLC)
SSTR2 Watson strand (LUAD)
SSTR2 Watson strand (SCLC)
SSTR2 Crick strand (LUAD)
SSTR2 Crick strand (SCLC)

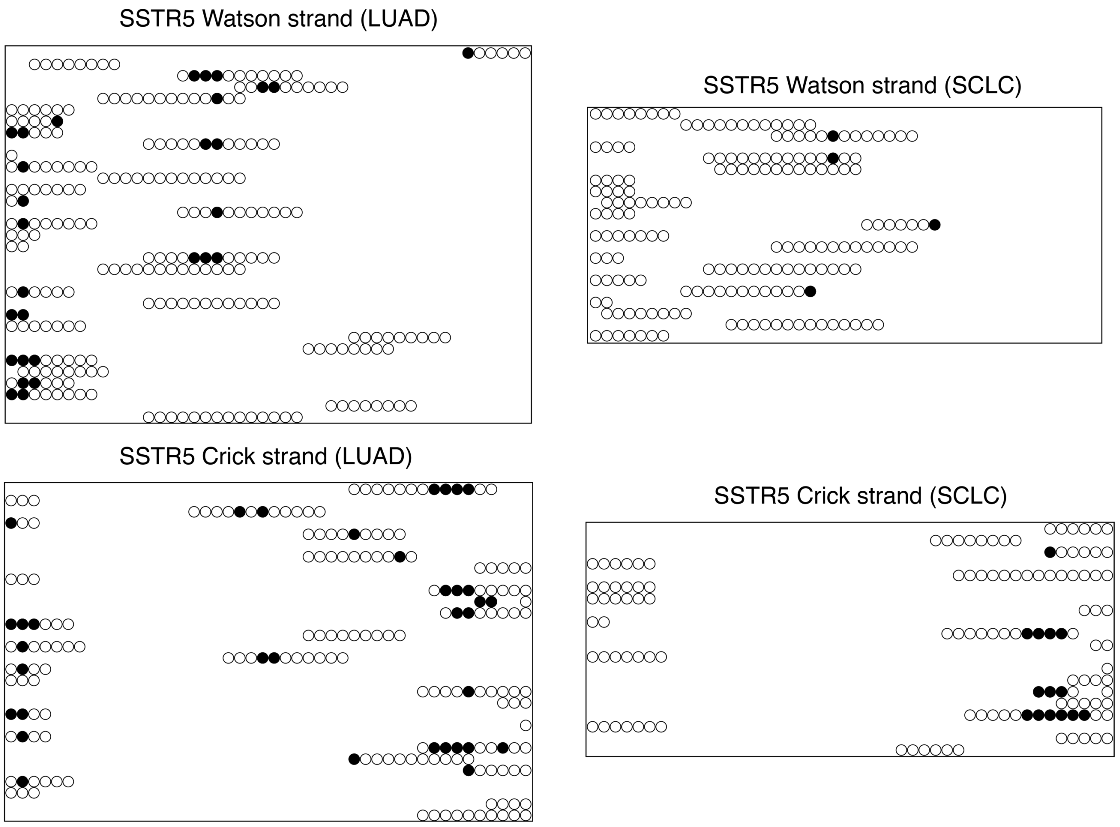
SSTR5 Watson strand (LUAD)
SSTR5 Watson strand (SCLC)
SSTR5 Crick strand (LUAD)
SSTR5 Crick strand (SCLC)

**b**

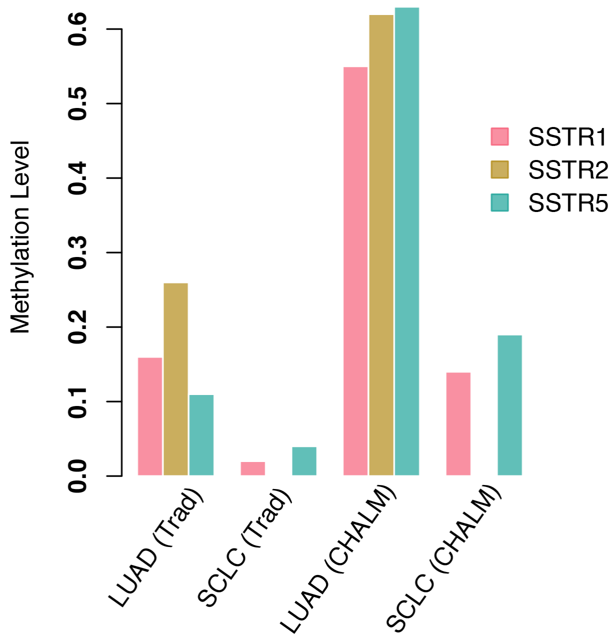
Methylation Level
0.0
0.1
0.2
0.3
0.4
0.5
0.6
SSTR1
SSTR2
SSTR5
LUAD (Trad)
SCLC (Trad)
LUAD (CHALM)
SCLC (CHALM)

**Supplementary Figure 27. Methylation status of the reads mapped to SSTRs. (a)** Raw sequencing reads mapped to the promoter regions of SSTR1, SSTR2 and SSTR5. For SSTR1 and SSTR2, the regions showed are the CHALM hypomethylated DMR between LUAD and SCLC comparison. For SSTR5, the region showed is the promoter CGI of this gene. Up to 50 reads are selected for each plot. Black circle: mCpG. White circle: CpG. **(b)** Grouped bar plot shows the traditional methylation and CHALM methylation level of three SSTR genes in LUAD or SCLC.

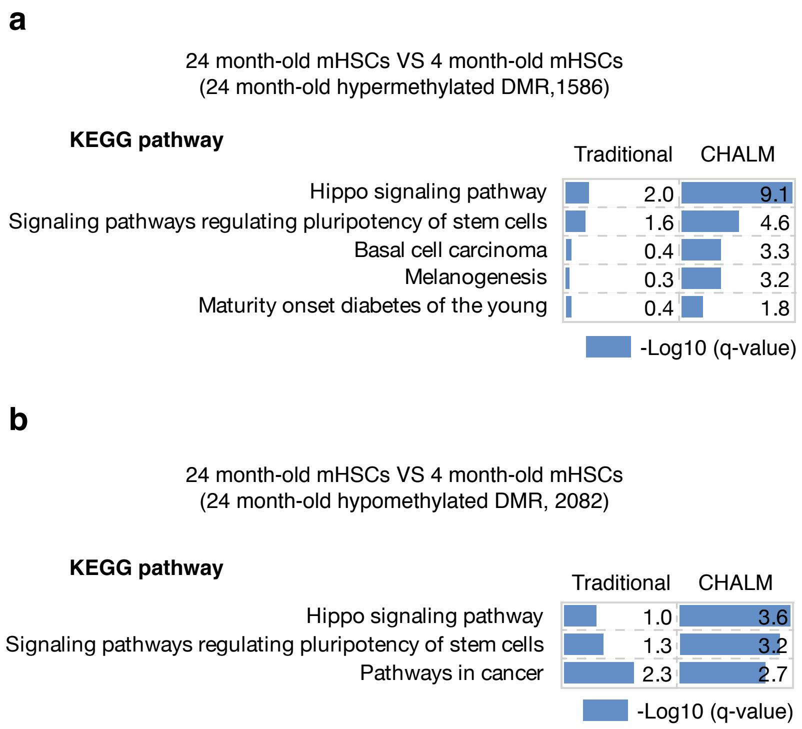


**Supplementary Figure 28. CHALM identified de novo DMRs that are more related to the underlying biological processes (old-young mHSC comparison).** The pathway enrichment analysis of the hypermethylated DMRs **(a)** and hypomethylated DMRs **(b)** in old mHSC.

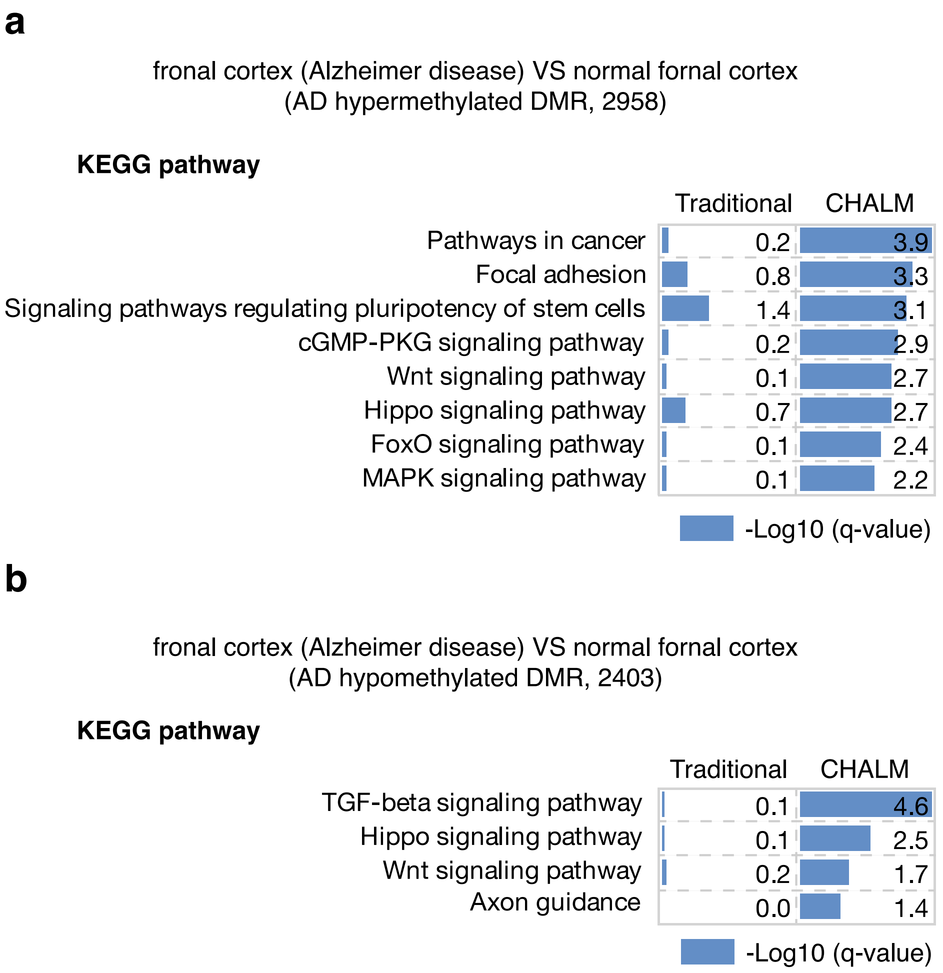


**Supplementary Figure 29. CHALM identified de novo DMRs that are more related to biological processes (AD-normal comparison).** The pathway enrichment analysis of the hypermethylated DMRs **(a)** and hypomethylated DMRs **(b)** in AD sample.

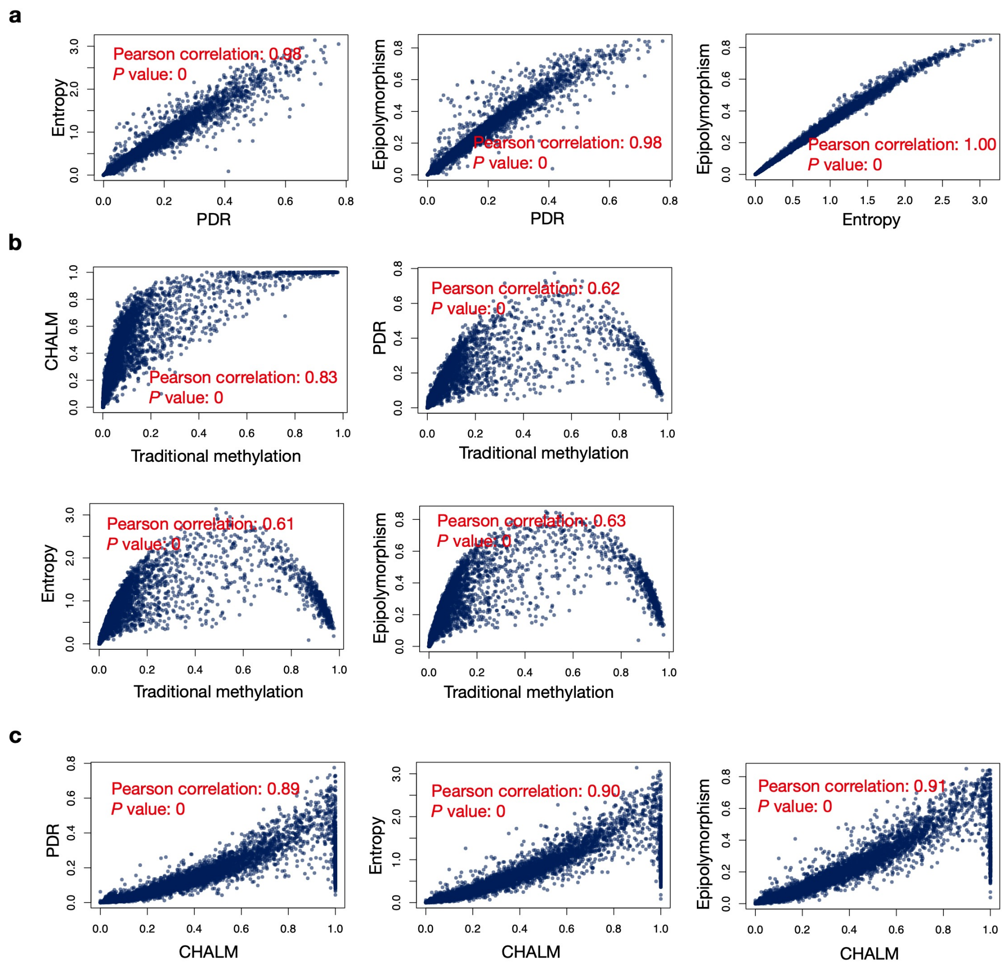


**Supplementary Figure 30. Comparison of CHALM to PDR, Epipolymorphism and Entropy with CD3 primary cell. (a)** Comparison between three heterogeneity methods. **(b)** Comparison of CHALM and the heterogeneity methods to the traditional mean methylation in promoter CGIs. **(c)** Scatter plots showing the comparisons of CHALM to heterogeneity methods.

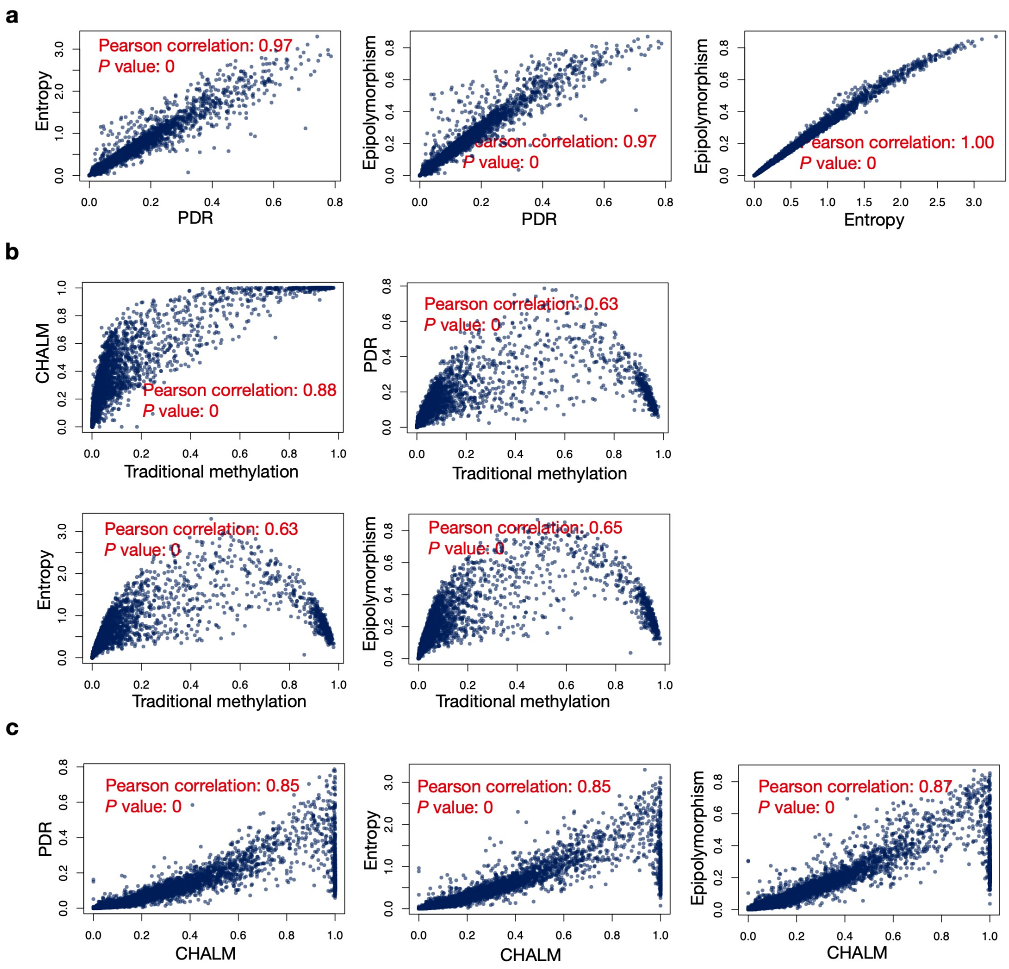


**Supplementary Figure 31. Comparison of CHALM to PDR, Epipolymorphism and Entropy with CD14 primary cell. (a)** Comparison between three heterogeneity methods. **(b)** Comparison of CHALM and the heterogeneity methods to the traditional mean methylation in promoter CGIs. **(c)** Scatter plots showing the comparisons of CHALM to heterogeneity methods.

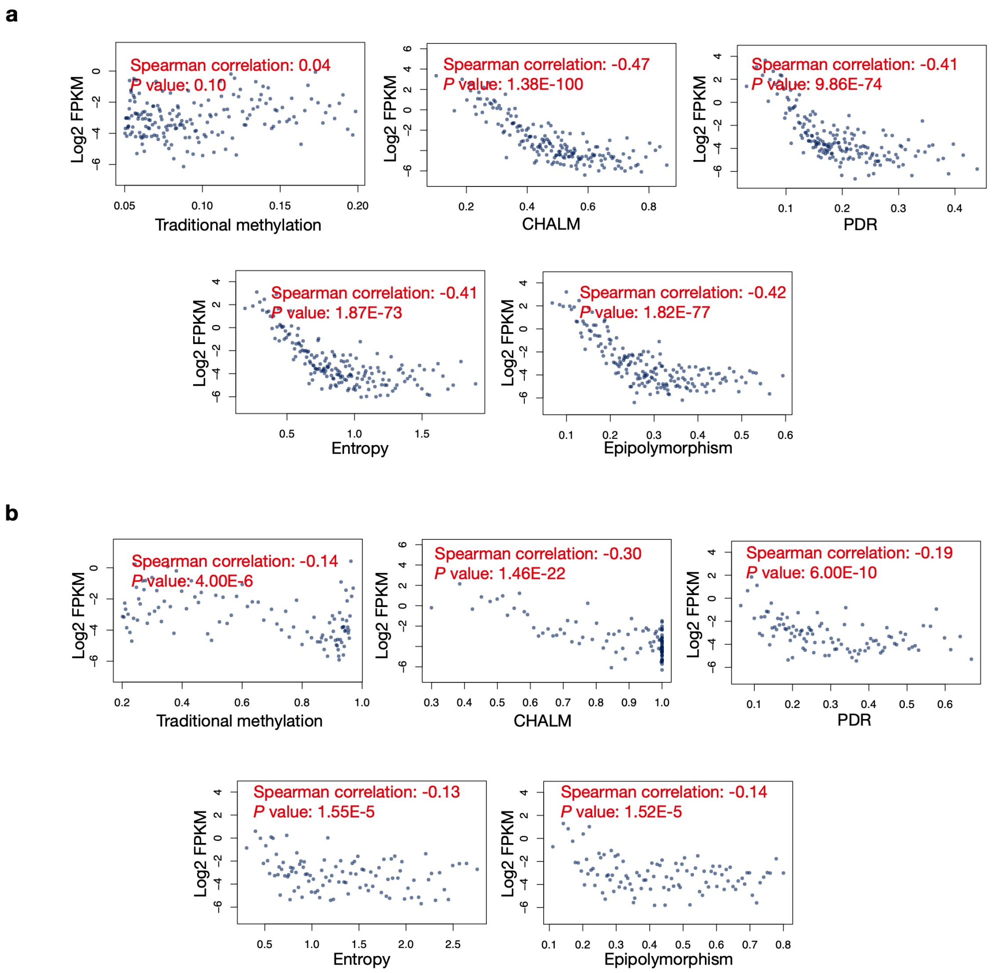


**Supplementary Figure 32. Correlation of heterogeneity methods with gene expression in the promoter CGIs of CD3 primary cell. (a)** Correlation of heterogeneity methods with gene expression for low methylated genes (traditional methylation between 0.05 and 0.2). **(b)** Correlation of heterogeneity methods with gene expression for high methylated genes (traditional methylation between 0.2 and 1).

a

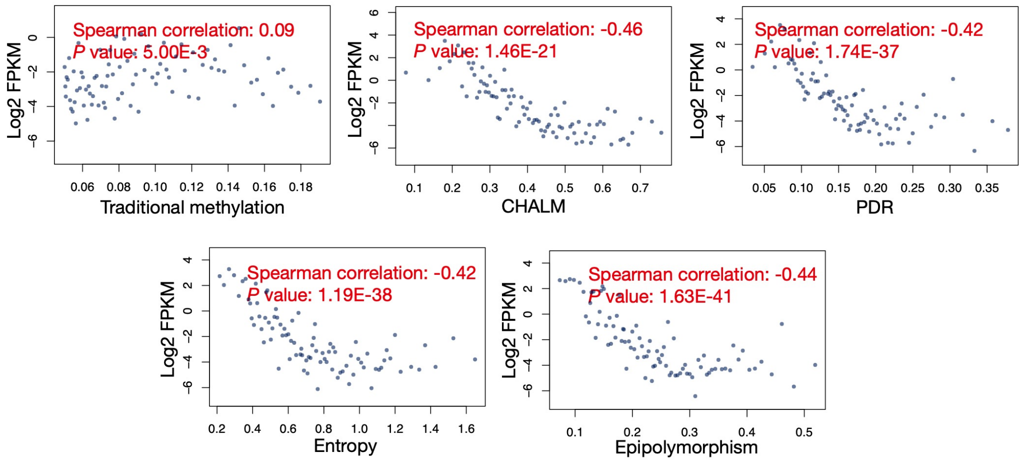


b

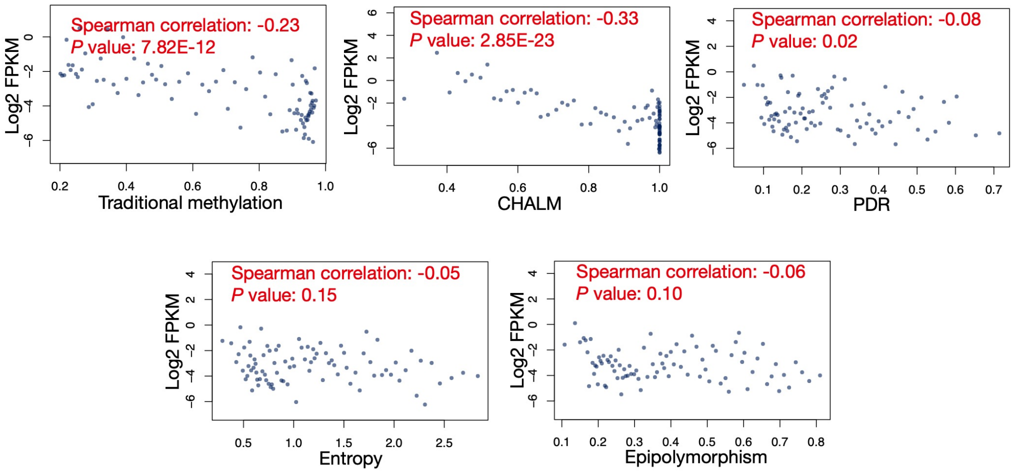


**Supplementary Figure 33. Correlation of heterogeneity methods with gene expression in the promoter CGIs of CD14 primary cell. (a)** Correlation of heterogeneity methods with gene expression for low methylated genes (traditional methylation between 0.05 and 0.2). **(b)** Correlation of heterogeneity methods with gene expression for high methylated genes (traditional methylation between 0.2 and 1).

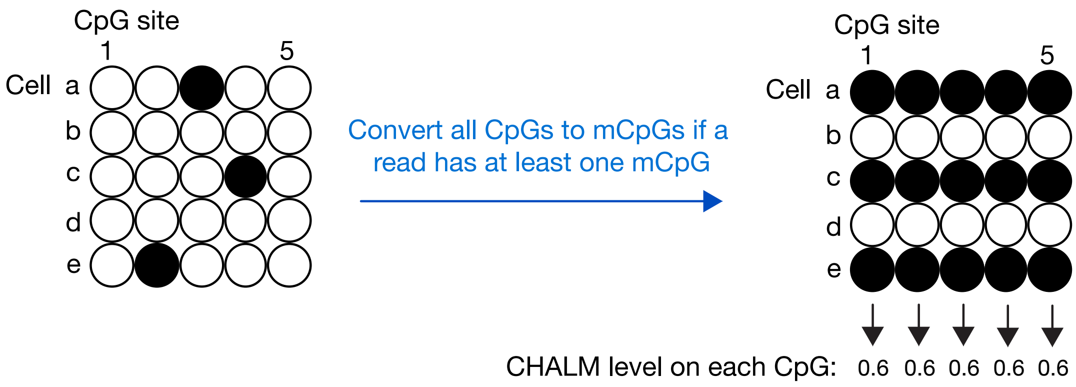


**Supplementary Figure 34. CHALM implementation.** First, all CpGs will be converted to mCpGs if a read has at least one mCpG. Then, CHALM level on each CpG site can be quantified as the ratio of mCpG. This implementation method of CHALM is compatible to most existing downstream tools, such as DMC and DMR calling tools.

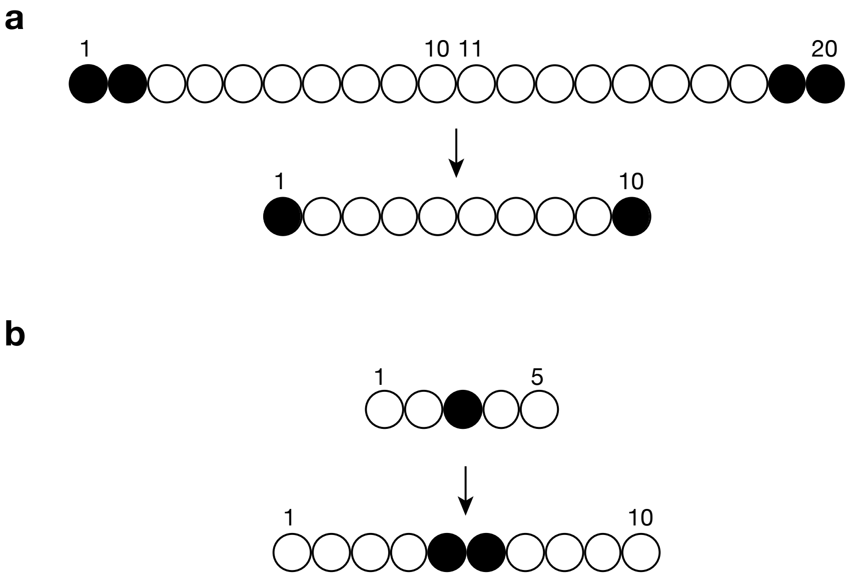


**Supplementary Figure 35. Process the raw sequencing reads into the Deep learning input matrix**. Each raw sequencing read is processed to be a row of the input matrix with the length of 10, which represents 10 CpG sites. (a) For the scenario that a read has $N$ $(N > 10)$ CpG sites, $Q$ and $R$ are the quotient and remainder of $N/10$. Then, for the first $(Q + 1) \times R$ CpG sites from left to right of this read, every $(Q + 1)$ CpG sites will be merged into a new CpG site by averaging, generating the first $R$ CpG sites. For the rest $Q \times (10 - R)$ CpG sites, every $Q$ CpG sites will be merged into a new CpG site, generating the rest $(10 - R)$ CpG sites. **(b)** For the scenario that a read has $N$ $( 5 \leq N < 10)$ CpG sites, $Q$ and $R$ are the quotient and remainder of $10/N$. Then, for the first $(10 - N)$ CpG sites of the read, every CpG site is duplicated into two adjacent CpG sites, generating the first $(20 - 2 \times N)$ CpG sites. For the rest $(2N - 10)$ CpG sites of this read, their methylation status will be copied to the rest CpG sites of the final row.
